# Ancestral and recent bursts of transposition shaped the massive genomes of plant pathogenic rust fungi

**DOI:** 10.1101/2025.01.10.632365

**Authors:** Emma Corre, Emmanuelle Morin, Sebastien Duplessis, Cecile Lorrain

## Abstract

**Background:** Transposable elements (TEs) play a crucial role in genome evolution, influencing gene regulation, diversity, and genome architecture. Rust fungi of the order Pucciniales (Subphylum Pucciniomycotina) are the largest group of obligate biotrophic plant pathogens and harbor some of the largest and most TE-rich genomes - up to 1.2Gb - compared to other fungi. This global genomic expansion contrasts with the smaller genomes and minimal mobilome found in other Pucciniomycotina species. Despite the availability of high-quality genome assemblies, our understanding of TE dynamics in Pucciniales remains limited due to inconsistent and incomplete TE annotations.

**Results:** We analyzed the mobilomes of 12 Pucciniomycotina species, producing a manually curated TE library for each genome. In Pucciniales, TEs occupy 47–92% of the genome, whereas 2-36% of TEs are detected in other Pucciniomycotina genomes. The comparison of gene and TE repertoires indicates that TEs, particularly LTR-retrotransposons and TIR-DNA transposons, are the primary contributors to the genome expansion of the Pucciniales. We reconstructed the proliferation histories of TEs in the Pucciniales, combining sequence similarity, clustering, and molecular clock approaches. We highlight recent and ancient TE invasions with some LTR-Gypsy elements predating the divergence of Pucciniomycotina (∼176 Mya), while most TE accumulation in Pucciniales occurred within the last 50 Mya. However, the TE invasions in the Pucciniales genomes do not seem to result from specific deficiencies in known TE-control mechanisms.

**Conclusion:** Our findings uncover extensive TE proliferation in Pucciniales, predominantly driven by LTR-Gypsy expansions. The retention of ancestral TEs and the consistently TE-rich genomes observed in Pucciniales highlight TE proliferation as an ancestral genomic feature in rust fungi.

## BACKGROUND

Rust fungi (order Pucciniales) are ubiquitous plant pathogens that infect a wide range of vascular plants, including some of the world’s most important crops and trees, such as wheat, soybean, poplar, and coffee trees [1, 2]. Epidemics caused by rust diseases can lead to significant yield losses, posing a substantial threat to global food security [2–4]. The Pucciniales are classified within the subphylum Pucciniomycotina, phylum Basidiomycota, in the Fungal Kingdom and represent the largest order of obligate biotrophic fungi, comprising over 8,000 species [5]. The Pucciniales are estimated to have emerged between 175-230 million years ago [5, 6]. Phylogenetic studies suggest that the diversification of rust fungi is closely tied to their obligate host association and coevolution, with occasional host jumps to unrelated species [7]. The obligate host association has shaped the genomic architecture of rust fungi. For instance, Pucciniales genomes show a reduced repertoire of plant cell wall-degrading enzymes, secondary metabolites, and transporter genes, typical traits of obligate biotrophic fungi [8–10]. However, Pucciniales genomes are exceptionally large, ranging from approximately 80 Mb to over 1.2 Gb [11, 12], and are expected to present even greater genome sizes [13], contrasting with the average ∼46.5 Mb genome size commonly found in Basidiomycota [14]. These expanded genomes are characterized by lineage-specific multigene families, an abundance of secreted proteins encoding genes, and a high proportion of repetitive elements [8, 9, 11]. The genome size expansion observed in Pucciniales parallels other filamentous plant pathogens. For instance, several Ascomycetes and Oomycetes, such as *Blumeria graminis* and *Phytophthora infestans*, also possess large genomes exceeding 100 Mb, similarly characterized by a significant proportion of repetitive elements [15]. The high repeat content of Pucciniales genomes has posed challenges in generating high-quality genome assemblies, which are crucial for studying their organization and evolutionary dynamics.

The increasing affordability of new-generation sequencing technologies has revolutionized the sequencing of high-quality reference genomes for numerous non-model organisms, including species within the Pucciniales order [11, 16, 17]. The most recent genome assemblies consist of gapless and phased chromosomes generated from dikaryotic spores [11, 18]. Analyses of the most recent Pucciniales genomes have confirmed that transposable elements (TEs) are major drivers of genome architecture and evolution in rust fungi [11, 17]. TEs are ubiquitous mobile DNA sequences capable of proliferating within their host genomes. TEs are classified into two main classes based on transposition mechanisms [19]. Class I elements, or retrotransposons, replicate through an RNA intermediate, enabling their insertion into new genomic positions. Class II elements, or DNA transposons, move either through direct excision from double-stranded DNA or via single-strand excision followed by a rolling-circle replication mechanism [19]. These two major classes are further organized into Orders and Superfamilies (SFs) based on their sequence architecture [19]. Within each class, elements are grouped into Families that share phylogenetic ancestry. Both classes include autonomous elements, which encode the enzymatic machinery necessary for their own mobilization, and non-autonomous elements, which depend on the transposition machinery provided by autonomous counterparts. In Class I TEs, autonomous elements include for instance long terminal repeat (LTR) elements and long interspersed nuclear elements (LINEs). In Class II TEs, autonomous elements primarily consist of terminal inverted repeat (TIR) elements and Helitrons [19]. Conversely, non-autonomous elements encompassing Miniature Inverted-repeat Transposable Elements (MITEs), Short Interspersed Nuclear Elements (SINEs), Terminal-repeat Retrotransposons in Miniature (TRIMs), and LARDs (Large Retrotransposon Derivatives), which lack the coding capacity for independent transposition [19].

TE-mediated genomic plasticity may confer significant adaptive advantages in plant-pathogen interactions. TEs contribute to genomic plasticity by facilitating chromosomal rearrangements, promoting compartmentalization, duplicating or deleting genes, and influencing gene expression [20]. In the soybean rust *Phakopsora pachyrhizi*, a subset of TEs is expressed alongside effectors during the early stages of infection, suggesting that transposition activity may be influenced by the plant-rust interaction, as also reported in other pathosystems [11, 21]. There is also evidence that fungal genomes have co-opted some TEs [22]. For instance, the effector AvrK1 in the TE-rich *B. graminis* f. sp. *hordei* genome is derived directly from a LINE retrotransposon [23]. The interplay between TE expansion and transposition regulation is largely unknown in TE-rich fungal species, including Pucciniales. However, these species appear to have evolved tolerant genomes that can accommodate TE activity, unlike other fungal species with compact genomes [24]. This tolerance may play a crucial role in the adaptability of rust fungi, contributing to their success as plant pathogens.

TE propagation can harm host genomes, and species have evolved defense mechanisms to regulate TE proliferation. In fungi, TE activity is often controlled through epigenetic silencing mechanisms [25, 26]. TEs are frequently associated with heterochromatin histone marks and cytosine DNA methylation (5mC) [27]. Repeat-induced point mutation (RIP) is also a defense mechanism that has been so far specifically found within Ascomycetes [8]. RIP inactivates TEs through cytosine-to-thymine mutations and involves the DNA methyltransferases DIM2 and RID [25, 27, 28]. To date, RIP has not been functionally characterized in Basidiomycetes, although RIP-like mutational signatures have been reported in the genomes of some smut fungi [29]. Notably, the canonical RIP-associated DNA methyltransferases, DIM2 and RID, are absent from the gene repertoires of multiple Basidiomycete species, including the Pucciniales [30]. RNA interference (RNAi) adds another layer of TE control. The RNAi pathway relies on three core components: Dicer, which processes double-stranded RNA into small interfering RNAs (siRNAs); Argonaute, which incorporates these siRNAs to guide the RNA-induced silencing complex (RISC); and RNA-dependent RNA polymerase (RdRP), which amplifies siRNAs to sustain the silencing response [31]. This pathway targets TE transcripts for degradation or chromatin modifications, thereby repressing TE expression and preventing transposition [32]. Despite its demonstrated importance in plants and animals, RNAi-mediated TE regulation in fungi remains comparatively less studied. Comparative genomic studies suggest that fungi exhibit significant diversity in their TE control mechanisms, with some species retaining these pathways while others have lost them through evolution [25, 27, 31].

The present study investigates the role of TEs in shaping Pucciniales genome architecture over evolutionary timescales. We tested four key hypotheses: (i) TE invasions in Pucciniales are the result of an ancestral transposition burst that has been preserved, (ii) TE invasions in Pucciniales arise from species-specific independent bursts, (iii) TE proliferation reflects a combination of ancestral bursts retained within the genome and recent species-specific transposition events, and (iv) TE proliferation is an ancestral characteristic of Pucciniomycotina evolution, which has been largely lost in other species but retained in Pucciniales. To achieve this, we generated a comprehensive annotation and curation of mobilomes across 12 selected Pucciniomycotina species with comparative genomic analyses, focusing on TE diversity, abundance, and evolutionary dynamics. Additionally, we compared gene content and patterns of multigene family expansion and contraction across Pucciniomycotina genomes to detect potential evidence of TE control presence/absence.

## RESULTS

### The massive genome size of Pucciniales is determined by their extensive mobilomes and large gene catalogs

To explore the contributors to the genome size expansion in Pucciniales, we selected a panel of 12 Pucciniomycotina species with reference genomes comprising eight Pucciniales species and four other Pucciniomycotina species (Fig. 1A; Table S1). We also included the complete genomes of two *Puccinia graminis* f. sp. *tritici* strains to compare intra-specific variation. The size of the nine selected Pucciniales genomes ranges from 109 Mb (*Melampsora larici-populina*) to 1.057 Gb (*P. pachyrhizi MT2006*), contrasting with the much smaller genomes of the four other Pucciniomycotina genomes, with sizes ranging from 20 Mb (*Rhodosporidium toruloides*) to 26 Mb (*Microbotryum lychnidis-dioicae*) (Table S1; Fig. 1B). Assemblies included either primary scaffolds from unphased builds or fully separated haplotype-phased scaffolds. BUSCO (basidiomycota_odb10) completeness was uniformly high (≤ 10 % missing for all, except *Leucosporidiella creatinivora* at 10.6 %), and most showed low duplication (≤ 10 % duplicated BUSCOs). However, the three Pucciniales assemblies *M. allii-populina* (22 %), *M. americana* (44 %) and *P. pachyrhizi* (36 %), exhibited elevated BUSCO duplication, consistent with uncollapsed haplotypes in these dikaryotic, repeat-rich genomes. To mitigate potential bias for comparisons, phylogenetic analyses were made on single-copy orthogroups and performed TE comparisons on a global genomic scale.

**Fig. 1:**
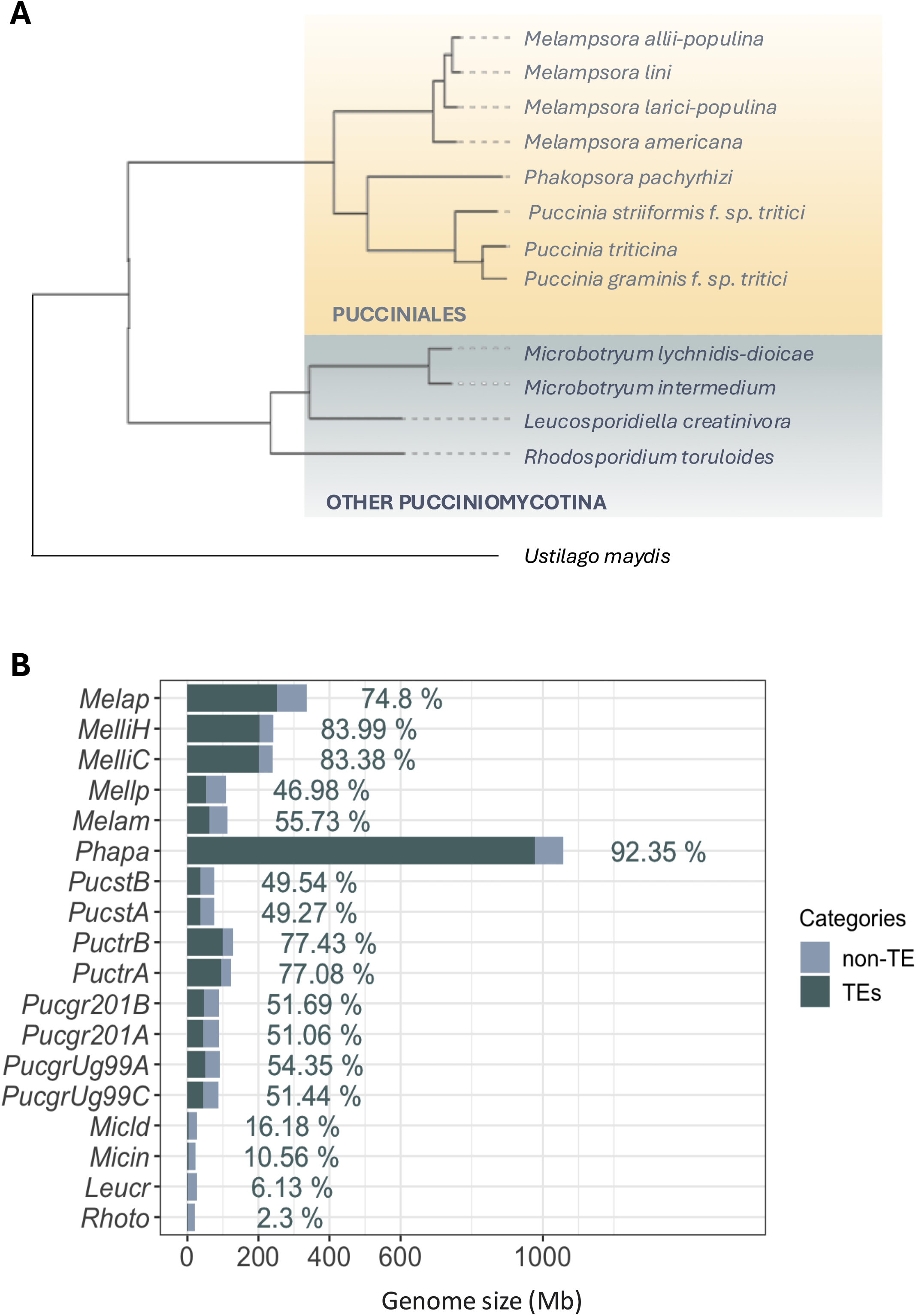
The phylogenetic tree and transposable element content in the Pucciniomycotina. A) phylogenetic tree of the 12 selected Pucciniomycotina species based on a single-copy orthogroup from Orthofinder v2.4.0 and visualized using iTOL (https://itol.embl.de/). The Pucciniales are represented in yellow. The category *Other Pucciniomycotina* (blue) refers to the four species of Pucciniomycotina outside the order Pucciniales. B) The bar plot displays the TE content across the analyzed genomes. Dark blue bars represent the proportion of the genome covered by TEs, while light blue bars show the total genome size (in Mb). Percentages on the right indicate the TE coverage relative to the total genome size. Species are described as follow: Melap (*Melampsora allii-populina*), Melli (*Melampsora lini*), Mellp (*Melampsora larici-populina*), Melam (*Melampsora americana*), Phapa (*Phakopsora pachyrhizi*), Pucst (*Puccinia striiformis* f. sp. *tritici*), Puctr (*Puccinia triticina*), PucgtrUg99 (*Puccinia graminis* f. sp. *tritici* Ug99), Pucgtr201 (*Puccinia graminis* f. sp. *tritici* 20-1), Micld (*Microbotryum lychnidis-dioicae*), Micin (*Microbotryum intermedium*), Leucr (*Leucosporidiella creatinivora*), Rhoto (*Rhodosporidium toruloides*).

We first compared gene and TE contents and their contribution to the genome inflation of Pucciniales. Pucciniales genomes harbor substantially more genes ranging from 15,984 genes in *Melampsora americana* to 19,618 genes in *P. pachyrhizi,* compared to the other Pucciniomycotina species with 7,819 genes in *M. lychnidis-dioicae* to 9,854 in *L. creatinivora* (Table S1). Although gene coverage showed a slight but significant positive relationship with genome size (r² = 0.42, p < 0.05), TE content appeared as the primary driver of genome inflation in Pucciniales, correlating very strongly with genome size (r² = 0.91, p < 0.001; FigS1). In addition, the TE coverage in Pucciniales genomes ranges from 47% to 92.3% in *M. larici-populina* and *P. pachyrhizi*, respectively (Fig. 1B; Table S2). We observed variation of the TE coverage within the Pucciniales order and between closely related species. For instance, in the Melampsoraceae, TE content represents 47.0% of the 109 Mb genome of *M. larici-populina* to 83.4-84% of the 239.9-242.3 Mb genome of *Melamspora lini C* and *M. lini H,* respectively. In the Pucciniaceae, TEs covered from 49.3-49.5% of the 75.6-75.9 Mb genome of *Puccinia striiformis* f. sp. *tritici* A and B, respectively, to 77.1-77.4% of the 123.9-121.6 Mb genome of *Puccinia triticina* A and B, respectively (Fig. 1B). In comparison, the TE coverage in the other Pucciniomycotina ranges between 2.3% in *R. toruloides* and 16.2% in *M. lychnidis-dioicae*. Finally, haplotype-phased Pucciniales assemblies display nearly identical TE proportions between phases, further validating our global-TE comparison approach. Together, these results highlight that TE accumulation, more than gene number, underlies the extensive genome expansion in Pucciniales relative to other Pucciniomycotina.

### LTR retrotransposons are the major contributors to the extensive mobilome and genome size expansion in the Pucciniales

To detect if TE proliferation in the Pucciniales is linked to specific elements, we further examined the TE landscapes in the genomes of the selected Pucciniomycotina species using REPET v3.0 followed by three rounds of manual curation to build a high-quality Pucciniomycotina TE library. In total, we annotated 39, 834 TE consensus sequences, constituting the Pucciniomycotina TE library. TE annotation before curation detected 20, 958 (52.6%) classified TE consensus sequences, 16,811 (42.2%) unclassified TE consensus sequences, and 2,065 (5.2%) TE consensus sequences with ambiguous annotations (or “conflicts”), which occur when the classifier cannot assign a TE to a single category due to nested or overlapping elements (Fig. S2). After the first round of manual curation, we reduced the number of ambiguous annotations to 651 (1.6%) TE consensus sequences but could not resolve unclassified annotations. Therefore, we used MCHelper [33] and improved the annotation of ∼16% (2, 700) of previously unclassified TE consensus sequences (Fig. S2). We used a final step during which we clustered all TE consensus sequences from the Pucciniomycotina TE library to reduce redundancy and correct the remaining ambiguous annotations. Our final curated Pucciniomycotina TE library comprised 28,868 (72.5%) classified TE consensus sequences and 10,966 (27.5%) unclassified TE consensus sequences (Tables S2-S4).

On average, the Pucciniales genomes are covered by 33.9% of Class I TEs, ranging from 15.4% (16.9 Mb) in *M. larici-populina* to 51.7% (515.8 Mb) in *P. pachyrhizi*, and 23.5% of Class II TEs ranging from 13.2% in *M. lini* haplotype C (31.6Mb) to 31.8% in *Puccinia triticina* haplotype A (39.3 Mb) (Fig. 2A; Table S2). By contrast, the Class I and Class II genome coverages were drastically lower within the other Pucciniomycotina, with an average of 13.7% and 3.5%, respectively (Fig. 2A; Table S2). Class I elements represent the most abundant elements in the Pucciniales except for the three genomes of *M. larici-populina* with 15.4% Class I (16.9 Mb), *Melampsora americana* with 18.8% Class I (21.1 Mb) and *P. striiformis* f. sp. *tritici* 18.1% Class I (13.7 Mb) (Fig. 2A; Table S2). These results indicate an important contribution of Class I elements in the genome expansions compared to the Class II elements in six of the eight Pucciniales species.

**Fig. 2:**
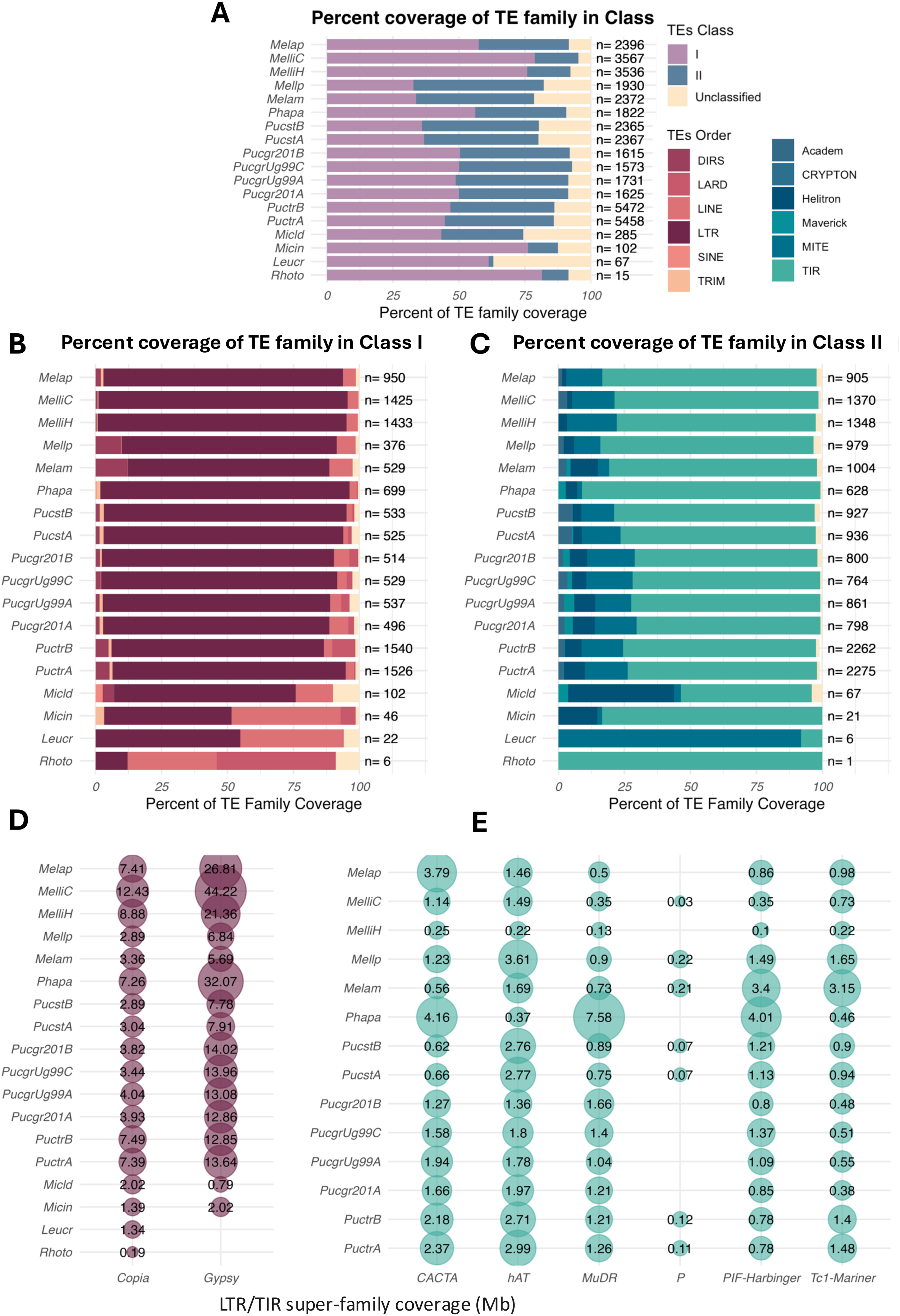
Composition of transposable elements in Pucciniomycotina genomes. The stacked bar plots represent the percentage of transposable elements (TEs) annotated at the A) Class level, B) Class I order level, and C) Class II order level for each Pucciniomycotina genome. The n values indicate the number of distinct TE families identified in each genome. The percentage of TE genome coverage at the super-family (SF) level for E) Class I elements (purple) and F) Class II elements (blue).

The TE Orders content of the Pucciniales genomes predominantly comprises LTR retrotransposons and TIR DNA transposons. LTRs represent 12.5% of the genome in *M. larici-populina* (13.1 Mb) to 63.0% in *M. lini* haplotype C (151.1 Mb), respectively (Fig. 2B-C; Table S3). At the LTR super-family level, we mostly found Copia and Gypsy elements, representing 2.9% of genome (3.2 Mb of LTR-Copia in *M. larici-populina*) to 12.7% (30.5 Mb in *M. lini* haplotype C) and 5.5% (6.2 Mb in *M. americana*) to 48.2% (115.6 Mb in *M. lini* haplotype C), respectively (Fig. 2 D-E; Table S4). The non-LTR retrotransposons, such as LINE elements, represent from 0.4% to 2.6% of the Pucciniales genomes. The non-autonomous elements such as SINE represent less than 0.2% of the Pucciniales genomes (Tables S3 and S4). In the other Pucciniomycotina genomes, the amount of LINE covers up to 3.3% of the genome of *M. intermedium* (0.77 Mb). TIRs represent most DNA transposons within the Pucciniales genomes, with 10.2% to 28.1%, followed by non-autonomous MITE representing 0.6% to 5.0% (Table S3). Interestingly, some TIRs super-family like CACTA, hAT, MuDR, PIF-harbinger, and Tc1-Mariner are found only in the Pucciniales and not in the other Pucciniomycotina (Fig. 2 D-E; Table S4). The Helitron elements cover a small proportion of the genome, from 0.2% in *M. lini* haplotype C (6.2 Mb) to 2.6% in *P. triticina* haplotype A (3.2 Mb). These results indicate that at the TE-Order level, the Pucciniales display an overall similar TE landscape with a majority of LTRs and TIR elements, suggesting ancient proliferation of TEs in the order Pucciniales. However, the variable amount of LTR and TIR elements found between Pucciniales genomes, even at the haplotype level, support additional bursts of transposition that occurred throughout the evolution of the order Pucciniales.

### Temporal dynamics of TE activity shaped the genome architecture of Pucciniales

We hypothesized that the proliferation of TEs observed in the Pucciniales could have emerged through different scenarios throughout the evolution of the Pucciniomycotina. First, we analyzed genome-wide TE divergence profiles using Kimura 2-parameter (K2P) substitution rates. In this framework, low K2P values signify recent TE insertions with minimal divergence from the consensus, whereas higher values denote ancient insertions indicative of long-term genomic retention and mutation accumulation. Our comparative landscape analysis across 12 Pucciniomycotina species revealed species-specific TE activity patterns (Fig. 3A–P). Among non-Pucciniales species, TE proliferation appears minimal. For instance, *R. toruloides* and *L. creatinivorum* exhibit low recent TE activity, with less than 3% of their genomic content composed of insertions with K2P < 0.05. Conversely, *M. intermedium* and *M. lychnidis-dioicae* demonstrate moderate levels of recent activity, with 8.5% and 10.5% of their genomes, respectively, reflecting young TE insertions (Fig. 3A–D). Within Pucciniales, two distinct TE proliferation patterns emerge. Species from the Melampsoraceae family show evidence of dual TE expansion waves with recent bursts (K2P < 0.05) accompanied by older insertions peaking between 0.4 and 0.6 K2P (Fig. 3E–I). Notably, *M. lini* presents a prominent accumulation of ancient LTR elements, indicating substantial historical activity. In contrast, members of the Pucciniaceae exhibit a predominance of recent insertions with a marked peak at K2P < 0.05 and a flatter distribution at higher divergence values, suggesting reduced retention of older TE sequences. In addition, we used a within genome sequence similarity-based approach to compare TEs from the oldest insertions to the most recent insertions as follows: i) divergent TE consensus sequences containing copies with less than 85% sequence identity, ii) intermediate TE consensus sequences with copies of 85% to 95% sequence similarity and iii) conserved TE consensus sequences comprising copies with more than 95% sequence identity [11, 12]. The average sequence identity of all TE consensus sequences per genome ranges from 80.6% (*R. toruloides*) to 90.6% (*M. lychnidis dioicae*). We observed 13.0--66.7% divergent (*M. lychnidis-dioicae* and *R. toruloides*, respectively), 33.3-76.5% intermediate (*R. toruloides* and *M. intermedium*, respectively), and 0.0-16.5% conserved (*R. toruloides* and *M. lychnidis-dioicae*, respectively) TE consensus sequences in the other Pucciniomycotina (Table S5). Within the Pucciniales, TE consensus sequences similarity profiles revealed 30.0-66.2% divergent (*P. pachyrhizi* and *M. larici-populina*, respectively), 27.4-65.6% intermediate (*M. americana* and *P. pachyrhizi*, respectively), and 2.7-7% conserved TE consensus sequences (*M. allii-populina* and *P. triticina*, respectively) (Table S6). Among the Pucciniales genomes, we observed that LTRs and TIRs compose most of each relative TE age category (Table S6). LTR elements represent 17.4-33.6% of the divergent TEs (in *M. americana* and *M. allii-populina*, respectively), while TIR elements represent 26.1-38.0% of the divergent TEs (in *P. pachyrhizi* and *M. larici-populina*, respectively). In the conserved TEs, LTR elements represent 7.2-34.4% (in *M. larici-populina* and *M. lini*, respectively), and TIR elements represent 7.6-33.3% (in *P. graminis* f. sp. *tritici Ug99 haplotype C* and *M. larici-populina*, respectively) (Table S6). The heterogeneity in the composition of TEs with different relative conservation profiles (i.e., relative insertion ages) in the Pucciniales suggests that multiple waves of TE proliferation occurred since the early divergence between the Pucciniales and the other Pucciniomycotina species. The subset of highly conserved elements in all genomes also indicates recent TE transposition events.

**Fig. 3:**
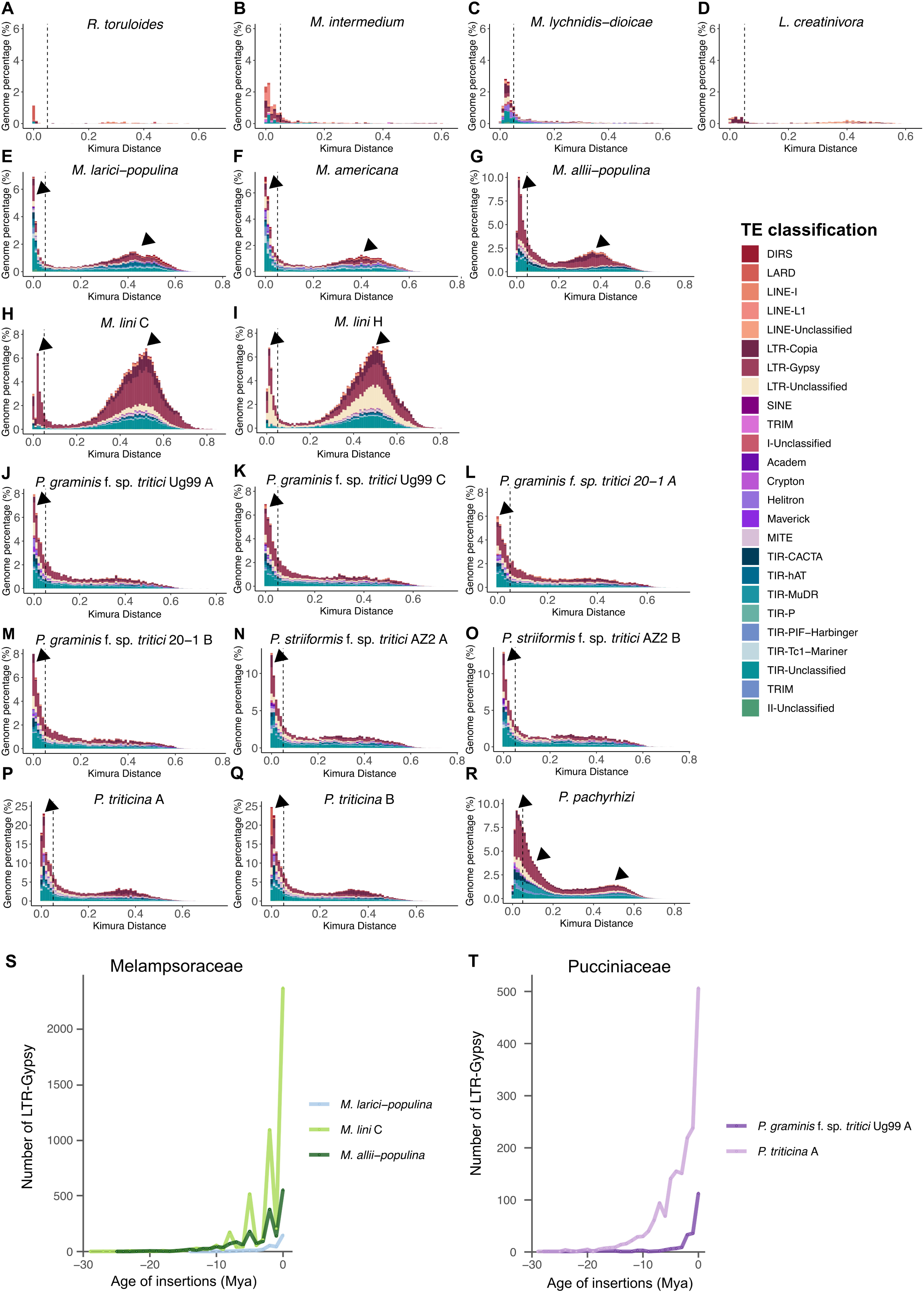
Transposable element insertion ages. A-R Kimura distances across the TE landscapes of Pucciniomycotina genomes. Each distribution shows the proportion of the genome (y-axis) composed of copies exhibiting varying degrees of divergence (x-axis). Arrows indicate TE mobilization events. S-T) Number of LTR-Gypsy elements accumulated through time (million years ago - Mya), based on insertion age for the S) Melampsoraceae species and T) the Pucciniaceae species.

Based on sequence similarity among all TE consensus sequences, we employed a clustering approach to identify TE families shared among or specific to species within the Pucciniales order. We found no TE family shared across all the Pucciniales species. However, we identified three clusters that are specific to the Melampsoraceae and one that are specific to the Pucciniaceae (Fig. S3). These findings suggest that the accumulation of ancient TE copies predated the divergence of the family Melampsoraceae and the family Pucciniastraceae (58±17 million years ago, according to Aime and McTaggart 2021 [2]) on one hand and the divergence of the family Pucciniaceae and the family Sphaerophragmiaceae (approximately 68±17 million years ago, according to Aime and McTaggart 2021 [2]) on the other hand. Interestingly, the three Melampsoraceae-specific TE clusters were all TIR elements, whereas the Pucciniaceae-specific clusters comprised one LTR cluster (Fig. S3; Tables S7 and S8). Additionally, we observed a predominance of species-specific TE clusters, ranging from 650 clusters (3.8% of the mobilome) in *P. graminis* f. sp. *tritici* 20-1 to 3,174 clusters (15.6% of the mobilome) in *P. triticina* (Fig. S3). Within these species-specific clusters, TIR elements accounted for 3.5% of the mobilome in *M. americana* (608 clusters) but only 0.05% (9 clusters) in *P. graminis* f. sp. *tritici* 21-0.

Interestingly, we identified haplotype-specific TE clusters in *M. lini* exhibiting 744 clusters (4.3% of its mobilome), whereas *P. graminis* f. sp. *tritici* showed between 42 clusters in Ug99 and 45 clusters in 20-1 haplotypes (Fig. S3). These findings highlight the variation in TE composition and specificity across species and even between haplotypes within individuals.

Finally, we retraced the evolutionary history of LTR invasion using their terminal repeats with Kimura molecular clock estimates. Across all Pucciniales genomes, LTRs were found to have accumulated over a broad temporal range, from 618 Mya to the present (Fig. S4). The oldest Gypsy elements were identified in *P. pachyrhizi*. 0.1% of LTRs accumulated before 176 Mya, thus predating the divergence between the Pucciniales and the other Pucciniomycotina (Fig. S4). However, our analysis suggests that LTR accumulation within the Pucciniales genomes, more particularly, spiked in the last 50 Mya, with a notable acceleration reflecting LTR expansions occurring within the last 5 Mya (Fig. 3S-T; Fig. S4). We also identified two distinct proliferation profiles in the Pucciniales genomes. The first profile, observed in *M. lini* and other Melampsoraceae species, exhibits sharp bursts of Gypsy LTR activity, with up to 2,000 elements accumulating during specific intervals (e.g., 7-9 Mya, 6-4 Mya, and 3-2 Mya) (Fig. 3S; Fig. S4). The second profile, characteristic of Phakopsoraceae and Pucciniaceae species, demonstrates a more gradual accumulation of LTRs over time (Fig. 3S-T; Fig. S4). These results collectively indicate that while genome inflation driven by TE invasion is a shared feature of Pucciniales, multiple waves of proliferation have shaped their evolutionary trajectories, reflecting diverse temporal dynamics of TE activity across lineages.

### The TE invasion of the Pucciniales appears unrelated to deficiency in TE control pathways

We investigated whether the TE invasion in Pucciniales compared to the other Pucciniomycotina, might be due to a loss of control mechanisms. The most studied control mechanism against TE proliferation in fungi is the induction of cytosine-to-thymine mutations in repetitive regions through RIP mutations [25]. Using the composite RIP index in TE sequences and in genome-wide 1-kb sliding windows, we quantified RIP-like signatures in TE family consensus sequences in the Pucciniales and the other Pucciniomycotina genomes. In all Pucciniomycotina genomes we detected less than 0.5% of the genomes containing RIP-like signatures (Table S9). We estimated the average RIP composite index per TE family ranging from −0.66 in *M. americana* to −0.15 in *L. creatinivora*, indicating the lack of RIP-like signatures in TE families among the Pucciniales and the other Pucciniomycotina TE family sequences (Table S9). Additionally, the two DNA methyltransferase genes associated with RIP (DIM2 and RID) are absent from both the Pucciniales and the other Pucciniomycotina genomes. We identified two other DNA methyltransferase genes DNMT5 and DNMT1 present across all Pucciniomycotina genomes, except for *P. hordei* in which we did not find DNMT1 (Fig. S5). Finally, we investigated the presence or absence of genes involved in the RNAi pathway, namely Argonaute, Dicer, and RNA-dependent RNA polymerase RdRP. We identified 25-72 (*P. hordei* and *P. triticina*) genes related to Dicer-like genes in the Pucciniales and 25-29 Dicer-like in the other Pucciniomycotina genomes (Fig. S6). Similarly, we identified 17-60 (and *M. lini*) RdRP-like genes in the Pucciniales and 12-15 RdRP-like genes in the other Pucciniomycotina genomes (Fig. S6). We observed a remarkable expansion of AGO1 solely in the *P. pachyrhizi* genome that displays 572 copies compared to a mean of 91 copies in the other species (Fig. S5). However, it remains to determine whether these copies are truly functional. All Pucciniomycotina genomes, regardless of their TE content, are equipped with genes potentially involved in TE control through DNA methylation silencing and RNAi pathways. We conclude that the TE proliferation observed in the Pucciniales genomes compared to the other Pucciniomycotina is not attributable to the loss of genes involved in TE control mechanisms and related to RIP, DNA methylation, or the RNAi pathway.

Finally, we analyzed gene family dynamics within the Pucciniomycotina subphylum to investigate potential evidence of loss of genes that may relate to TE control mechanisms in Pucciniales outside of the canonical mechanisms investigated above. For this, we determined multigene families within the Pucciniomycotina using *Ustilago maydis* as an outgroup and identified 43,847 orthogroups (Table S10). Of these, 3067 core orthogroups (17,1758 genes) were shared across all genomes, including *U. maydis*, while 16, 585 orthogroups (85,786 genes) were specific to Pucciniomycotina (Table S10). We observed substantial variation in species-specific orthogroups ranging from 68 in *M. intermedium* to 1310 in *P. triticina.* We identified 23 orthogroups in contraction and 380 orthogroups in expansion at the separation of the Pucciniales and the other Pucciniomycotina (Fig S7). The orthogroups in contraction in the Pucciniales comprise a total of 483 genes, of which 19.5% (94 genes) are of unknown functions (Table S11; Fig. S8). The five most represented functions within the remaining orthogroups in contraction in the Pucciniales, include 62 genes related to “Transcription”, 52 genes related to “Signal transduction mechanisms”, 52 genes with function related to “Amino acid transport and metabolism”, 25 genes related to “Posttranslational modification, protein turnover, chaperones”, 23 genes related to ““Secondary metabolites biosynthesis, transport and catabolism” and 21 genes related to “Chromatin structure and dynamics”(Fig S8; Table S11). However, we identified only 22 orthogroups with a significant accelerated gene gain/loss between the Pucciniales and the other Pucciniomycotina genomes and all the 22 orthogroups are in expansion in the Pucciniales (p-value < 0.05; Fig S9). These results show that Pucciniales underwent extensive gene losses compared to other Pucciniomycotina species. Based on the functional categories identified in this comparative analysis, it remains possible that certain gene losses may be associated with the loss of functions related to TE control within the ‘chromatin structure and dynamics’ category or other, as yet unidentified, functions.

## DISCUSSION

In this study, we provide a better understanding of how TEs have profoundly shaped the genomes of Pucciniales, contributing to their distinctive genome expansions within the Pucciniomycotina subphylum. Our comprehensive TE annotation across 12 Pucciniomycotina genomes demonstrates that the Pucciniales possess a mobilome landscape enriched in LTR Gypsy and TIR elements, a feature contrasting sharply with the smaller genomes and the minimal TE contribution in other Pucciniomycotina species. Our analyses examined four hypotheses on TE proliferation in Pucciniales and evidenced both ancient and recent TE bursts, especially from Gypsy elements, aligning with the third hypothesis: TE proliferation is driven by a combination of ancestral transposition bursts retained in the genome and recent species-specific independent transposition events throughout Pucciniales evolution. Hence, we uncover ancient TE proliferation likely predating the early divergent branch of the order Pucciniales, followed by further species-specific expansions. In addition, despite the extensive TE presence, our findings indicate no notable deficiency in known TE regulatory mechanisms in the Pucciniales genomes compared to the other Pucciniomycotina genomes, suggesting that Pucciniales may have rather evolved tolerance to TE proliferation or deployed other mechanisms to accommodate their abundant presence in the genome.

The accumulation of TEs in Pucciniales is an ancestral genomic trait. Pucciniales is a monophyletic group of species [5], all possessing inflated genomes enriched in TEs. We identified retrotransposon copies retained in Pucciniales genomes that predate this divergence, suggesting a long-term persistence of ancestral TEs within rust fungal genomes [5]. This TE retention pattern is consistent with observations in previous works of rust fungal genomes, which often harbor highly divergent, long-standing TE families [17]. Interestingly, we did not find any TE families shared among all *Pucciniales* species in our panel, suggesting that ancestral copies have diverged substantially over time. However, we detected TE families consistently retained at the genus level, including one (LTR-Copia) within the family *Pucciniaceae* and three others (TIR-Unknown, TIR-hat, and TIR-Tc1Mariner) within the family *Melampsoraceae*. The genus-specific conservation of certain TEs suggests that ancestral elements have been retained in distinct subgroups, which mirrors patterns observed in other fungal species. For example, a recent analysis of transposable element repertoires across 24 arbuscular mycorrhizal fungal genomes revealed the persistence of ancient TEs over long evolutionary periods [34]. Likewise, in large, complex plant genomes such as wheat and maize, retrotransposons are retained across extended timescales [35, 36]. The long-term retention of TEs across such diverse fungal and plant lineages suggests a common pattern of TE evolutionary dynamics, underlying their potential role in shaping their host-genome architecture.

*Gypsy* retrotransposons represent the largest proportion of TE content across Pucciniales genomes, followed by *Copia* retrotransposons and TIR elements. This abundance of retrotransposons is a common feature in other TE-rich fungal genomes, such as those of *B. graminis* and *Entomophthoraceae* spp. [37, 38]. We hypothesize that ancient *Gypsy* elements found in the order Pucciniales may have served as progenitors for more recent lineage-specific TE expansions observed across the Pucciniales genomes. Indeed, the dynamics of *Gypsy* element accumulation display extreme variation among Pucciniales species. For example, we observed a burst accumulation pattern in *M. lini*, marked by a rapid increase in *Gypsy* copies over a few million years. In contrast, species such as *P. pachyrhizi* exhibit a more continuous pattern of *Gypsy* accumulations. We also detected haplotype-specific TE families in the phased genomes of the two species *M. lini* and *P. graminis* f. sp. *tritici*, indicating that TE dynamics vary not only between species but also within the two haplotypes in dikaryotic Pucciniales. These differences suggest that *Gypsy* elements may undergo distinct cycles of activity in each species, with some elements remaining active and driving ongoing TE expansion while others persist as inactive remnants within the genome. Notably, the presence of still active elements aligns with the observed expression of certain TEs during host infection in *P. pachyrhizi* [11], further supporting the hypothesis that active TEs may still be involved in ongoing genomic changes during pathogen-host interactions.

TE richness is a shared genomic feature among different groups of fungi, which appears as a trait associated with obligate parasitic and symbiotic lifestyles. Similarly to the Pucciniales, other obligate biotrophic plant pathogens, such as *B. graminis*, the causal agent of powdery mildew in cereals, are characterized by a high proportion of TEs [39]. Enrichment in TE could enable rapid evolution through genomic plasticity in response to fast-evolving host immune systems and changing environmental conditions [24]. Outside of plant pathogens, obligate insect parasites within the fungal order Entomophthorales exhibit TE-rich genomes [38]. TEs are reported as the main drivers for genome expansion in this clade. TE-rich genomes extend beyond obligate parasites and represent a key feature of beneficial plant symbionts. Arbuscular mycorrhizal fungi and ectomycorrhizal fungi exhibit substantial TE content and expansions, although the role of these TEs in their symbiotic functions remains scarcely understood [34, 40]. While the evolutionary pressures differ between parasitism and symbiosis, large TE-rich genomes in both contexts may suggest that TEs could reflect an evolutionary strategy to facilitate genome plasticity and adaptation to the host environment.

We observed no evidence of specific gene loss associated with known TE regulation mechanisms in the Pucciniales genomes compared to other Pucciniomycotina. First, RIP-like signatures are absent in the Pucciniomycotina subphylum which aligns with other studies centered on basidiomycetes and with the absence of DIM2 and RID genes in basidiomycetes [27, 41]. In addition, we found the DNA methyltransferases DNMT1 and DNMT5 in all 12 genomes, suggesting that the whole Pucciniomycotina phylum may possess the potential for TE silencing mediated by DNA methylation. DNMT1 and DNMT5 could play a role in the establishment and/or maintenance of cytosine DNA methylation (5mC), as found in plants and animals [27]. In addition, Bewick et al. (2019) [27] demonstrated that species in the Phylum Basidiomycota displayed the highest 5mC levels across fungal genomes, with a weak correlation between 5mC levels and repeat content in the selected species. Similarly, we identified the essential genes of the RNAi pathway in all 12 Pucciniomycotina genomes. Additionally, we confirmed the expansion of Argonaute 1 (AGO1) previously reported in *P. pachyrhizi* [11]. Most of these expanded AGO1-like genes were described as pseudo-genes surrounded by TEs, suggesting a TE-driven inactivation of expanded AGO1 gene members in the family. A comparable phenomenon occurs in the arbuscular mycorrhizal fungus *Gigaspora margarita*, where multiple AGO-like copies lack conserved PIWI catalytic residues and may have diverged to non-canonical roles or become pseudogenes [42]. This pattern could be similar to the recent loss of DIM2 activity found in the wheat fungal pathogen *Zymoseptoria tritici* where an expansion of the DIM2 genes, followed by inactivation, is thought to have led to TE-derepression [43]. In *Rhizophagus irregularis*, several AGO loci localize adjacent to TE insertions and transcriptional profiling has shown that AGO genes are highly expressed in spores but downregulated during plant colonization, coincident with transient TE activation [44, 45]. As our comparative genomic approach could not identify a Pucciniales-specific loss of TE control, other approaches, such as epigenomics or small RNA sequencing, will be required to better understand TE regulation mechanisms in rust fungi.

Despite advancements in TE annotation and the availability of high-quality rust genomes, the absence of a standardized and comprehensive TE library for rust fungi remains a significant obstacle in performing comparative genomics to address further evolutionary studies [46]. Manually curating TE libraries for highly repeated genomes is labor-intensive and time-consuming. Here, we generated a three-step approach to curate TE annotations, which can be applied to many other genomes. This curation has resolved all the ambiguous annotations and reduced the unclassified TEs from 42.4% to 16% after three rounds of manual curation. We created a unified rust fungal TE library, enabling consistent and precise TE identification across different rust fungal species. Still, several TE annotations remain unresolved after our curation, mainly consisting of elements probably too degraded to be correctly classified and nested elements. Additional manual curation efforts, supported by additional genome assemblies from closely related species, could support the refinement of the classification of these elements by offering more diagnostic features. As large-scale genomic projects generate vast sequencing data, creating a unified TE library becomes increasingly crucial for advancing our understanding of the roles of TEs in rust fungal biology and their evolution. Expanding and refining fungal TE libraries is essential to fully capture the diversity and dynamics of transposable elements in these genomes, enabling more precise evolutionary and functional insights in future studies.

## CONCLUSION

This study describes manually curated TE catalogs in thirteen Pucciniomycotina species and provides a new fungal TE database. We report the genome expansions in Pucciniales are mainly guided by TE proliferation. Our work highlights recent bursts of Gypsy LTR in Pucciniales while the content in such elements remains relatively poor in other Pucciniomycotina. We also describe a few conserved TEs within the Pucciniales order linked to older proliferation events. TE accumulations do not seem to result from Pucciniales-specific loss of TE control mechanisms. We provided new insights into Pucciniales genome architecture and favored the hypothesis that TE proliferation is an ancestral characteristic of the Pucciniales evolution.

## METHODS

### Genomic data

We selected the genomes and gene annotations of 12 Pucciniomycotina species publicly available in the JGI MycoCosm database and NCBI repositories (Table S1). We included ten genomes from the order Pucciniales, comprising four genomes from the family Melampsoraceae (*M. allii-populina*, *M. lini*, *M. larici-populina*, and *M. americana*), four genomes from the family Pucciniaceae (*P. graminis* f. sp. *tritici Ug99 and 21-0*, *P. triticina*, and *P. striiformis* f. sp. *tritici*), and one genome from the family Phakopsoraceae (*P. pachyrhizi*). Additionally, four genomes representing species from outside the order Pucciniales but within the subphylum Pucciniomycotina were included (*L. creatinivora*, *M. lychnidis-dioicae*, *M. intermedium*, and *R. toruloides*; Table S1). We selected genomes with either haplotype-phased assemblies, long-read assemblies, or assemblies based on genetic maps. We assessed genome assembly completeness using BUSCO v5.4.7 with the basidiomycota_odb10 marker set, which provided a measure of the quality of the selected genome assemblies (Table S1).

### TE detection and annotation

We detected and annotated repeats in Pucciniomycotina genomes using REPET v3.0 (https://urgi.versailles.inra.fr/Tools/REPET, [33]). Repeated elements were identified and annotated for each genome following a two-step process. First, TE families were identified using the TEdenovo pipeline, which builds TE family sequences based on repetitive elements detected in a 100 Mb subset of the genome. This subset-based approach was applied to enhance workflow efficiency in highly repeated genomes, as described in [34]. Subsequently, the whole genomes were annotated with TEannot using the TE families generated by TEdenovo. A second round of TEannot was performed as a curation step, focusing exclusively on TE families with at least one full-length copy. For haplotype-phased genome assemblies, TEdenovo was run separately for each haplotype (also using 100 Mb subsets), and the resulting TE family sequences were merged before proceeding with the TEannot steps. This approach allowed for the identification of potential haplotype-specific TEs.

### Curation of TE annotations

After the automated detection and annotation of TEs, we performed additional steps of curations of libraries to improve the classifications of elements detected as “conflicts” and “unclassified” TEs by REPETv3.0. First, the “conflicts” ambiguous TE annotations (i.e. the classifier cannot assign a TE to a single category) were manually curated and reassigned to the appropriate TE Class or Order based on sequence similarity, applying a 50% similarity threshold. In parallel, we used the Manual Curator Helper tool MCHelper [33] to address redundancy, fragmentation, false positives, and unclassified sequences in TE libraries for each genome. Briefly, MCHelper extends consensus sequences through iterative rounds of Blast, extract, and extend (BEE) and refines them into subfamilies based on distance clustering. False positives are filtered out by removing sequences with low-quality fragments or homology to multi-copy genes, and the remaining sequences are classified through homology-based methods. We then pooled all TE libraries into a Pucciniomycotina TE library to perform an additional manual curation step by clustering the Pucciniomycotina TE library with CD-HIT v4.8.1 [47]. We grouped TE sequences with 80% similarity across 80% of the sequence length and re-assigned previously unclassified TEs based on annotated TEs of the same cluster. The workflow to generate curated TE annotations from multiple genomes combining automated detection and curation results is available at https://github.com/Raistrawby/TE_Rust/tree/main. The curated library of TE consensus sequences has been deposited in the Dfam database (https://dfam.org/home).

### TE relative insertion age

The relative age of TE insertions in the Pucciniomycotina genomes was estimated with the similarity-based approach of REPETv3.0, as described in Lorrain et al. (2021) [48]. Briefly, we analyzed sequence divergence between TE copies to their cognate family using the extent of sequence similarity as a proxy to the divergence time of the TE copies within the host genome. The TE copies were classified into three categories: i) conserved, with copies similar up to 95% against their family, ii) intermediate, with a similarity between 85% and 95% and, iii) divergent, with similarity of copies below 85%. In addition, we estimated the sequence divergence of TE copies relative to their consensus sequences, we computed Kimura 2-parameter distances. For each annotated TE copy, the aligned region was extracted and compared to its corresponding consensus sequence. K2P distances were calculated based on the frequency of transitions and transversions and correcting for multiple substitutions following the approach described by Baril et al. (2024)[49] and adapted to our TE annotations (https://github.com/Raistrawby/TE_Rust/tree/main). Briefly, for each TE copy, the corresponding genomic sequence was retrieved based on genomic coordinates and strand orientation. Each extracted sequence was then pairwise aligned to its cognate consensus sequence using matcher [50] and the Kimura 2-parameter distance [51] was computed from the alignment. TE copies with incomplete or failed alignments were excluded from downstream analyses. The resulting Kimura distances were used as a proxy for the relative age of TE insertions.

### Insertion age of LTR-retrotransposons

The insertion ages of LTR-retrotransposons were assessed using a molecular clock approach, as described in Gupta et al. (2023) [11]. Full-length LTR-retrotransposons were detected using LTRHarvest v 1.6.1 [52] with default parameters (except for - similar 90 (default 85) and -mintsd 5 (default 4)). Gypsy and Copia LTR-retrotransposons were selected based on the annotations from the repbase_23.12 database. Each element’s 3’ and 5’ long-terminal repeats were extracted and aligned using MAFFT v7.471 [53] to calculate Kimura’s P distances [51]. Gypsy and Copia insertion dates (T) were estimated using the formula T_=_K / 2r, where K is the distance between the 2 LTRs and is calculated with the Kimura 2-Parameter distance formula −0.5*log(1 - 2p - q) * sqrt (1 - 2 q)) with p = transition frequency and q transversion frequency and r is the fungal substitution rate of 1.05 × 10-9 nucleotides per site per year [51].

### Gene family annotations and evolution

We predicted orthologs between the Pucciniomycotina genomes and the reference *U. maydis* genome [23], a representative of the Ustilaginomycotina (the closest subphylum to Pucciniomycotina within Basidiomycete), which we used as an outgroup to identify genes restricted to the Pucciniomycotina subphylum. Orthogroup analysis was performed using the software OrthoFinder v2.4.0 (i=2 - M MSA -S diamond -a mafft)[54] on the proteomes. We chose one haplotypic proteome for species with haplotype-resolved annotation. To assess the gene family gains and losses throughout the Pucciniomycotina genomes, we first needed to build a phylogenetic tree; to this end, we used the single copy orthogroups (i.e. orthogroups containing only one protein of each genome) and infer the tree with *U. maydis* as an outgroup with Raxml and 500 bootstraps [55]. We used this rooted phylogenetic tree and the number of genes found per family (i.e. orthogroup) to estimate rates of gene families gains (i.e., families in expansion) and losses (i.e. families in contraction) at the species-, genus-, family- and order-levels with CAFE v5.0 [56].

### Functional gene annotations and identification of DNA methyltransferases and RNA-induced silencing complex

We used EggNOG-mapper v2.1.12 [57] to predict functional annotations for all Pucciniomycotina genomes. We retrieved the PFAM, KOG, KEGG, and GO annotations from EggNOG-mapper annotations to explore the potential functions of gene families gained or lost throughout the evolution of the Pucciniomycotina. To identify the DNMTs and the genes involved in the RISC pathway in the Pucciniomycotina genomes, we extracted the genes with corresponding PFAM domains. For DNA methyltransferases, we selected genes with PF00145 (C-5 cytosine-specific DNA methylase) and used orthogroups and BLASTp searches to differentiate DNMT classes. For the RNA-induced silencing complex, we selected genes with PF05183 (RNA dependent RNA polymerase), PF16486 (N-terminal domain of argonaute), PF22389 (Protein Dicer, dsRNA-binding domain), PF09692 (Argonaute siRNA chaperone (ARC) complex subunit Arb1), PF22749 (Arb2 domain), PF00636 (Ribonuclease III domain), PF04992 (RNA polymerase Rpb1, domain6), PF03871 (RNA polymerase Rpb5, N-terminal domain), PF01191 (RNA polymerase Rpb5, C-terminal domain). We searched for characterized DNMT and RISC proteins to refine the annotations and performed BLASTp searches on the putative Pucciniomycotina DNMT and RISC proteins.

### RIP signatures analysis

RIP-like signatures were searched by estimating the RIP composite index [25] and GC content in TE family sequences of all Pucciniomycotina genomes, as described in Feurtey et al. (2023)[58]. RIP Composite index values were calculated using the formula: (TpA/ApT) – (CpA + TpG/ApC + GpT) applied per TE family sequence. A region is considered affected by RIP if the Composite index exceeds 0 [41]. In addition, we performed a 1kb genome-wide sliding window RIP-like signature detection using TheRIPper [59].

## Data availability

The accession numbers and links to publicly available genome assemblies and gene annotations used in this study can be retrieved from Table S1. The TE annotations and gene functional predictions generated in this study, are openly available at https://doi.org/10.5281/zenodo.14616359.

## Code availability

The TE annotation and manual curation pipeline, as well as additional scripts used to create the results presented in this manuscript, are available in the following GitHub repository: https://github.com/Raistrawby/TE_Rust/tree/main.

## Authors contributions

EC and CL wrote the manuscript. SD and EM revised the manuscript. EC, CL, and SD designed, analyzed, and interpreted the data. EM provided comparative genomics expertise and bioinformatics technical support. CL and SD contributed to the conception of the work.

## Acknowledgments

Jana Sperschneider and Peter Dodds are acknowledged for providing early access to the phased assembly and genome annotation of the *M. lini* reference genome.

## AI language model assistance

ChatGPT (developed by OpenAI) was used to refine and improve the clarity and grammar of the text. The authors reviewed and revised all outputs to ensure accuracy and alignment with the intended message.

## ETHICS DECLARATIONS

### Ethics approval and consent to participate

Not applicable.

### Consent for publication

Not applicable.

---

**Fig. S1:** Association between genome size and gene/TE content. Genome sizes (Mb) were plotted against genes or TE coverage for each Pucciniomycotina species. The lines represent the linear regression.

**Fig. S2: Improvement of TE annotations through manual curation**. The proportion of annotated TE families after REPET detection and annotation (RawR), after the first round of curation (Postpct), after the second round of curation with MCHelper (postMC), after a third round of curation with Pucciniomycotina-guided clustering (PostCDhit).

**Fig. S3: Shared and specific TE families between the Pucciniomycotina species.** Numbers of TE families shared among the 14 genomes based on CDHIT clustering. The bars on the left represent the total number of genes per species. The clusters shared among Pucciniaceae are colored in blue and the clusters shared among Melampsoraceae are colored in red.

**Fig. S4:** Number of LTR-Gypsy elements accumulated through time (million years ago - Mya), based on insertion age for each genome.

**Fig. S5: TE controls involved in methylations are detected in all species**. The number of genes involved in TE control corresponds to DNMT1 and DNMT5.

**Fig. S6: TE controls involved in RNA-induced silencing complex are detected for all genomes.** The number of genes involved in TE control corresponds to the RNAi pathway: Dicer-like (DCL; DCR), RNA-dependent RNA polymerase (RDP; RNT; RPAB; RPB), Argonaute (AGO; ARB) for each genome.

**Fig. S7:** Contraction and Expansion of multigene families in Pucciniomycotina. The phylogenetic tree of the selected Pucciniomycotina species based on single-copy orthogroups from Orthofinder v2.4.0 and infer with Raxml (500 bootstraps) and visualized using iTOL, rooted with *Ustilago maydis.* Numbers in purple correspond to the number of orthogroups in contractions at each branch or node and yellow at expansions.

**Fig. S8: Contraction of Multigene Families in Pucciniales.** Functional annotation of multigene families in contraction in Pucciniales. Genes are re-annotated by EggNOG-mapper v2.1.12. A threshold is set to kept only species with at least 20 proteins per COG category. COG categories correspond to: “-“: no hit; B: chromatin structure and dynamics; C: energy production and conversion; E: amino acid transport and metabolism; G: carbohydrate transport and metabolism; I: lipid transport and metabolism; K: transcription; O: posttranslational modification, protein turnover, chaperones, P: inorganic ion transport and metabolism; Q: secondary metabolites biosynthesis, transport and catabolism; S: function unknown; T: signal transduction mechanisms.

**Fig. S9: Significant contraction and expansion of multigene families in Pucciniomycotina.** A) The phylogenetic tree of the selected Pucciniomycotina species based on a single-copy orthogroup from Orthofinder v2.4.0 and visualized using iTOL, rooted with *Ustilago maydis.* Numbers in black correspond to the number of significant orthogroups in expansion/contraction at each branch or node. B) COG category annotation of genes involved in expansion and contraction between Pucciniales and other Pucciniomycotina. Contraction and expansion are detected with CAFE v5.0 and orthogroups with a *p*-value (0.05) are kept. Genes are re-annotated by EggNOG-mapper v2.1.12. The number of genes per species and categories. COG categories functions correspond to: A: RNA processing and modification; C: energy production and conversion; D: cell cycle control, cell division chromosome partitioning; G: carbohydrate transport and metabolism; I: lipid transport and metabolism; K: transcription; L: replication recombination and repair; M: cell wall, membrane, envelope biogenesis; N: cell motility; O: posttranslational modification, protein turnover, chaperones, Q: secondary metabolites biosynthesis, transport and catabolism; S: function unknown; T: signal transduction mechanisms.

**Table S1: General information of selected genomes** [60–70].

**Table S2: TE annotations at the Class level for the Pucciniomycotina genomes**. ClassAnn represents the different TE Classes (I, II, Unclassified); nbTE represents the number of TE families in each Class; ncovP represents the percentage of genome coverage of each TE class in the species genome; nbcopie represents the number of copies associated with the TE family for each Class; Sumcov represents the percentage of coverage by all the TEs per genome.

**Table S3: TE annotations at the Order level for the Pucciniomycotina genomes**. OrderAnn represents the different TE Orders; nbTE represents the number of TE families in each Order; ncovP represents the percentage of genome coverage of each TE Order in the species genome; nbcopie represents the number of copies associated with the TE family for each Order; Sumcov represents the percentage of coverage by all the TEs per genome.

**Table S4: TE annotations at the Super-family level for the Pucciniomycotina genomes**. OrderAnn represents the different TE SF; nbTE represents the number of TE families in each SF; ncovP represents the percentage of genome coverage of each TE SF in the species genome; nbcopie represents the number of copies associated with the TE family for each SF; Sommecov represents the percentage of coverage by all the TEs per species.

**Table S5: Average of the TE sequence identity at the Class level**. The meanID represents the mean sequence identity (%) of individual copies to their respective TE family consensus sequence averaged per TE class (ClassAnn).

**Table S6: Average of the TE sequence identity at the Order level**. The meanID represents the mean sequence identity (%) of individual copies to their respective TE family consensus sequence averaged per TE Order (OrderAnn).

**Table S7:** Shared TEs within the three Melampsoraceae genomes.

**Table S8: Shared TEs within the five Pucciniaceae genomes**.

**Table S9: RIP signature analysis**. Genome-wide RIP signature detection using TheRIPper [59]. RIP-like signatures were searched by estimating the RIP composite index in TE families consensus sequences for each Pucciniomycotina genome described in Feurtey et al. (2023)[58].

**Table S10: Orthogroups summary metrics of the selected Pucciniomycotina genomes.** Genomes are described as follow: Melap (*Melampsora allii-populina*), Melli (*Melampsora lini*), Mellp (*Melampsora larici-populina*), Melam (*Melampsora americana*), Phapa (*Phakopsora pachyrhizi*), Pucst (*Puccinia striiformis* f. sp. *tritici*), Puctr (*Puccinia triticina*), PucgtrUg99 (*Puccinia graminis* f. sp. *tritici* Ug99), Micld (*Microbotryum lychnidis-dioicae*), Micin (*Microbotryum intermedium*), Leucr (*Leucosporidiella creatinivora*), Rhoto (*Rhodosporidium toruloides*).

**Table S11:** Orthogroups in contraction of the selected Pucciniomycotina genomes.

